# Syn-TEF1 restores the functional disease phenotype of SCA27B in a hiPSC-derived neuronal model

**DOI:** 10.64898/2025.12.24.696231

**Authors:** Felix Gsell, David Pellerin, Oleg Vinogradov, Stephan L. Zuchner, Carolin Haag, Ulrike B.S. Hedrich, Marek Napierala, Mario Saporta, Christopher Yanick, Bernard Brais, Mara Baumgartner, Holger Lerche, Niklas Schwarz, Matthis Synofzik

**Affiliations:** Department of Neurology and Epileptology, Hertie-Institute for Clinical Brain Research and Hertie Center of Neurology, University of Tübingen, Tübingen, Germany; Dr John T. Macdonald Foundation Department of Human Genetics and John P. Hussman Institute for Human Genomics, University of Miami Miller School of Medicine, Miami, USA; Department of Neurology, University of Miami Miller School of Medicine, Miami, Florida, USA; Division Translational Genomics of Neurodegenerative Diseases, Hertie Institute for Clinical Brain Research and Hertie Center of Neurology, University of Tübingen, Tübingen, Germany; German Center for Neurodegenerative Diseases (DZNE), University of Tübingen, Tübingen, Germany; Department of Neurology, Peter O’Donnell Jr. Brain Institute, UT Southwestern Medical School, Dallas, USA; Department of Neurology and Neurosurgery, Montreal Neurological Institute, McGill University, Montreal, Quebec, Canada

**Keywords:** SCA27B, FGF14, iPSC-derived neurons, Na^+^ voltage-gated channel, action potential, neuronal excitability, transcriptional elongation

## Abstract

Spinocerebellar Ataxia 27B (SCA27B), caused by a deep-intronic GAA•TTC repeat expansion in the first intron of the *FGF14* gene, is one of the most frequent genetic ataxias. Its underlying disease mechanisms remain largely unknown, and disease-modifying therapies targeting upstream processes are lacking. Here we hypothesized that (i) SCA27B is driven by transcriptional repression of *FGF14*, which encodes a protein regulating ion channels at the axon initial segment (AIS), resulting in reduced Na^+^ channel availability and neuronal excitability, and that (ii) these defects can be restored by a synthetic elongation transcription factor (Syn-TEF1).

We assessed *FGF14* mRNA levels by qPCR and neuronal function by whole-cell patch-clamp recordings in iPSC-derived neurons from two SCA27B patients and two healthy controls. Patients carried GAA•TTC repeat expansions that were either monoallelic (391/16 repeats) or biallelic (315/290 repeats), exceeding the common pathogenicity threshold of >250 repeats. *FGF14* mRNA levels were reduced approximately to 60% and 70% of control levels in monoallelic and biallelic SCA27B neurons, respectively. This was accompanied by impaired excitability, with cumulative action potential (AP) firing reduced to 38% and 45% of control levels in monoallelic and biallelic lines, respectively, and peak Na^+^ current density reduced to 46% and 41%, while voltage-dependent gating of Na^+^ channels remained unchanged. Treatment with Syn-TEF1 significantly increased *FGF14* mRNA expression and restored cumulative AP firing to 83% and 135% of control levels in monoallelic and biallelic neurons, respectively, and Na^+^ peak current density to 95% and 138%.

These findings strongly suggest that the pathophysiological cascade in SCA27B – from *FGF14* repression to impaired Na^+^ currents and decreased neuronal excitability – can be reversed by an elongation transcription factor. Our results thus provide a rationale for further exploring Syn-TEF1 as a first gene-targeted, disease-modifying therapeutic approach for SCA27B.

## Introduction

Spinocerebellar Ataxia 27B (SCA27B) has recently emerged as one of the most frequent genetic forms of cerebellar ataxia in many populations worldwide [1, 2]. Clinically, it manifests as a late-onset, slowly progressive neurodegenerative disorder characterized by (vestibulo-) cerebellar motor symptoms, such as gait and limb ataxia, and downbeat nystagmus. Episodic ataxia often presents early in the disease course, and major involvement of other neurological systems or cognitive impairment are uncommon [3, 4]. SCA27B is caused by a deep intronic GAA•TTC repeat expansion in the first intron of the Fibroblast Growth Factor 14 (FGF14) encoding gene (*FGF14*), with the TTC triplet located on the sense strand. While the disease is commonly caused by a single repeat expansion on one allele (monoallelic), expanded repeats on both alleles (biallelic) are also frequent [5, 6]. Despite its high frequency and substantial disease burden, the pathophysiological mechanisms resulting from *FGF14* repeat expansions in SCA27B remain poorly understood, and therapies targeting these upstream disease mechanisms remain to be developed.

The FGF14 protein itself is critically involved in modulating the function of ion channels and neuronal excitability [1, 5, 7–19]. It is a member of the intracellular FGF family, also known as Fibroblast Growth Factor Homologous Factor 4 [10, 20–22], and binds directly to ion channels (Na_V_1.2, Na_V_1.6 and K_V_7) as well as to the cytoskeletal scaffold protein ankyrin G [14, 15, 23]. Through this dual interaction, FGF14 ensures the precise spatial alignment and anchoring of ion channels at the AIS, which is essential for the initiation of action potentials (AP) and modulation of firing frequency and neuronal excitability [7, 10, 13, 24].

In contrast to other autosomal-dominant repeat expansion disorders that often act through toxic RNA or protein gain-of-function mechanisms [25], SCA27B appears to follow a different molecular-cellular pathophysiology. The pathogenic GAA•TTC repeat expansion, located in the first intron of the *FGF14-1b* isoform, causes a significant reduction in its transcript levels [1]. Reduced FGF14 expression is hypothesized to impair vol tage-gated ion channel function, which might lead to deficits in AP generation [7, 19]. However, there is so far no direct evidence linking this transcriptional deficiency to functional neuronal deficits in a SCA27B cellular model.

Emerging insights into gene regulation reveal that expanded GAA•TTC repeats can cause transcriptional blocks through RNA polymerase II (Pol II) pausing during elongation, which prevents full-length mRNA production [26–30]. Productive elongation requires release from this paused state, a step regulated by elongation factors such as BRD4 and P-TEFb [31]. Building on this mechanism, synthetic transcription elongation factors (Syn-TEFs) have been developed to specifically target genomic loci with repeat expansions containing guanine and adenine, and recruit the endogenous transcription machinery, enabling Pol II to overcome repressive chromatin or structural barriers imposed by repeat expansions [26, 32–34]. For example, Syn-TEF1 was shown to restore expression of frataxin in cardiomyocytes and sensory neurons derived from induced pluripotent stem cells (iPSCs) derived from patients with Friedreich’s ataxia. By specifically binding to expanded GAA•TTC repeats, Syn-TEF1 selectively recruits elongation complexes, enabling locus-specific transcriptional reactivation without effects on cells with unexpanded GAA•TTC repeats [26].

Here, we investigated the hypotheses that the GAA•TTC repeat expansion in *FGF14* similarly causes transcriptional elongation defects that underlie reduced FGF14 expression in SCA27B thereby impairing neuronal excitability. Using human induced pluripotent stem cell (hiPSC)-derived neurons from patients, we examined the impact of *FGF14* GAA•TTC on voltage-gated Na^+^ currents and AP firing and dynamics. Furthermore, we tested whether Syn-TEF1 could restore FGF14 expression and rescue potential functional deficits in SCA27B. Our findings aim to provide (i) the first mechanistic link between transcriptional dysregulation and neuronal dysfunction in SCA27B; and (ii) proof-of-concept for targeted transcriptional reactivation as a promising upstream therapeutic strategy for this recurrent disease.

## Materials and Methods

### Generation of human-induced pluripotent stem cells

Two hiPSC-lines from SCA27B patients, one with a monoallelic (B6440: 16/391) and one with a biallelic (B4796: 290/315) expanded GAA•TTC *FGF14* repeat expansion were generated from fibroblasts obtained from skin biopsies [1]. Two hiPSC-clones from healthy individuals (PIZ and WCMT2) with normal GAA•TTC *FGF14* repeat length were used as controls [35, 36] (number of GAA repeats determined as published previously [37]).

### Neurogenin2 integration

Stable Neurogenin2 (NGN2)-inducible hiPSC lines were generated via CRISPR/Cas9-mediated targeting of the AAVS1 safe harbor locus. Cells were nucleofected with two plasmids: pXAT2 (encoding Cas9 and AAVS1-targeting sgRNA) and pUCM-AAVS1-TO-hNGN2 (carrying NGN2, rtTA, and a puromycin resistance cassette flanked by AAVS1 homology arms). The pXAT2 and pUCM-AAVS1-TO-hNGN2 plasmids were gifts from Knut Woltjen (Kyoto University, Japan) and Michael Ward (University of Texas at Arlington, USA), respectively [38, 39]. Nucleofection was performed using the Nucleofector™ 2b device and Human Stem Cell Nucleofector™ Kit 2. Prior to transfection, cells were pre-treated with 10 µM Y-27632 (ROCK inhibitor). Post-nucleofection, cells were plated onto Matrigel-coated wells in mTeSR medium containing ROCK inhibitor. Puromycin selection (0.5 µg/mL for 72 h) began three days later. Successfully targeted clones were expanded and cryopreserved for further use.

### *NGN2*-based neuronal differentiation

Frozen stocks were prepared in cryopreservation media and stored in liquid nitrogen at - 196°C. Defrosted stocks of 500 µl were solved in 15 ml DMEM (12491015, Thermo Fisher) and centrifugated at 300g for 3 minutes. Cells were then seeded in mTeSR Medium (85850, StemCell Technologies) on matrigel-coated cover slips. For expansion, manually picked hiPSC colonies were cultured on matrigel-coated dishes (354277, Corning) with mTeSR Medium without antibiotics, changed daily, and passaged every 7 days. Only cells with passage numbers below 30 were used in experiments. Mycoplasma was not detected in any of the cell lines by PCR (G238, abm). HiPSCs were differentiated into cortical glutamatergic neurons by transient overexpression of the transcription factor Neurogenin 2 (NGN2) under the control of a doxycycline-inducible cassette. For differentiation into neurons, hiPSC cultures were grown on Matrigel until reaching 50–70% confluence. NGN2 expression was rapidly initiated, driving the cells toward a neuronal fate by adding 2.5μg/ml doxycycline (D9891, Sigma-Aldrich) for 5 hours followed by dissociation by accutase (A6964, Sigma-Aldrich) and plating 60,000 cells per coverslip on a 24-well plate in neuronal stem cell medium (NSC - 47.5% Neurobasal Medium (Thermo Fisher Scientific (Gibco), USA, 21103049), 47.5% DMEM/F12 (Thermo Fisher Scientific (Gibco), USA, 21331-020), 1% N2 supplement (N2 supplement Capricorn, Germany, N2-K), 2% NCS supplement (Capricorn, Germany, C21-H), 1% 100x GlutaMAX (Thermo Fisher Scientific (Gibco), USA, 35050087), 1% Non-essential amino acids (PAN-Biotech GmbH, Germany, P08-32100) containing 1 µg/ml doxycyclin, 10 µM Thiazovivin, and 3 µM of the notch pathway inhibitor DAPT (208255-80-5, StemCell technologies)). The following day, the medium was replaced with NSCM containing 10 µM DAPT, 10 µM PD0325901 (4192, Tocris), and 1 µg/mL doxycycline. After 1 week, 2 µM Cytosin-β-D-arabinofuranosid (Ara-C; C1768, Sigma-Aldrich) was added for at least 24h to get rid of remaining hiPSCs. After 8 days of differentiation, half of the medium was changed twice a week with pure NSC medium.

### Treatment of neuronal cultures with Syn-TEF1

Syn-TEF1 (PA1-PEG -JQ1) was synthesized from Pharmaron and used as described in [26]. Syn-TEF1 was administered two days before RNA extraction or at least three days before electrophysiological recordings. Coverslips (CSs) grown in wells with 0.5 ml (24-well plate, continental cultures) or 1 ml (12-well plate, autapses) of NSC medium were treated with either 0.5 or 1 µl of 1 mM Syn-TEF1 or 0.5/1 µl NSC only (control condition).

### Quantitative PCR on human-induced neurons

Total RNA was extracted from hiPSC-derived neurons cultured for two weeks without astrocytes on PLO/laminin-coated wells using the ISOLATE II RNA Mini Kit (BIO-52072, Meridian Bioscience). RNA concentration was measured with a NanoDrop ND-1000 (Peqlab, Germany). Quantitative PCR was performed in quadruplicates using PowerUp SYBR Green Master Mix (A25741, Applied Biosystems) with 10 ng RNA per reaction. FGF14 transcript levels were amplified using primers targeting the 1B isoform (Forward: 5′-CATATGCTGCAGTGTCTTTGTG-3′; Reverse: 5′-GTAGTAGCCTTGCCTGCAATA-3′), normalized to GAPDH, and quantified relative to the respective control or untreated condition using the ΔΔCt method.

### Electrophysiological recordings

Electrophysiological experiments were performed between DIV 30-37. Whole cell patch clamp recordings were performed at room temperature (22-24°C). For all experiments, the same extracellular (in mM: 140 NaCl, 4.2 KCl, 1.1 CaCl_2_, 1.0 MgSO_4_, 0.5 Na_2_HPO4, 0.45 NaH_2_PO_4_, 5 HEPES, 10 glucose pH: 7.4, Osmolarity: 305 mOsm) and intracellular solutions (in mM: 140 K-gluconate, 1 CaCl_2_, 10 EGTA, 2 MgCl_2_, 4 Na-ATP, 10 HEPES, pH 7.2, Osmolarity: 295 mOsm) were used. The electrophysiology rig was equipped with an Axopatch200B (Molecular Devices, USA) amplifier connected to a Digidata 1440A (Molecular Devices, USA) and running pClamp10 (Molecular Devices, USA) acquisition software. Voltage clamp recordings of Na^+^ currents were sampled at 100 kHz and filtered with 10 kHz, and cells were clamped at -70 mV. Series resistance (R_s_) was typically in the range of 5–10 MΩ and was compensated by ∼60–80%. Recordings were excluded if R_s_ exceeded 10 MΩ or changed by more than 20% during the experiment or holding current exceeded -100 pA. The estimated voltage error was monitored and remained below ∼10–20 mV for the currents analyzed. To minimize space clamp artefacts, recordings were restricted to neurons with simple morphology and small soma size. Only cells with stable access resistance and without signs of poor voltage control were included in the analysis. Steady-state inactivation of voltage-gated Na^+^ channels (VGSCs) was assessed using a voltage-clamp protocol consisting of 100□ms conditioning pulses to membrane potentials ranging from –90□mV to 0□mV in 10□mV increments, followed by a 10□ms test pulse to –10□mV. The holding potential was set to –70□mV throughout the protocol. To assess activation properties, voltage steps from –60□mV to +20□mV were applied in 10□mV increments (20□ms duration each), from a holding potential of –70□mV.

For isolating the M-current, voltage-gated Na^+^ and Ca^2+^ channels were inhibited by bath application of 1 µM tetrodotoxin and 0.5 mM CdCl_2_, respectively. Additionally, 5 mM 4-aminopyridine was used to block K_V_1 and K_V_3 channels. After a conditioning pulse at -10 mV, which eliminates the fast-activating and -inactivating K_V_1 and K_V_3 channels (A-current), voltage steps from -60 to +60 mV in 20 mV increments were used (duration: 500 ms). M-currents were defined as the non-inactivating component of outward potassium current sensitive to 20 μM XE-991 estimated from the last 100 ms. The protocol was regularly applied during the application of XE-991 to monitor the magnitude of XE-991 inhibition. Recordings of the XE-991-resistant current were only undertaken once there were no further changes in current amplitude. The current amplitude was normalized to each cell’s capacitance to obtain a measure of current density (pA/pF). In autaptic cultures, synaptic responses were triggered by 2 ms depolarization steps to 0 mV from a holding potential of 70 mV.

For the current clamp experiments, the sampling rate was 100 kHz and low pass filtered with 10 kHz. The holding current was set to achieve a membrane potential of −70 mV correcting for the liquid junction potential (15 mV, calculated from internal/external solutions using LJPcalc). The evoked firing was recorded using a protocol with 800□ms depolarizing steps in 25□pA increments from –25□pA to to 125□pA.

### Data analysis

Analysis of electrophysiological data was performed using Microsoft Excel for Microsoft 365 (version 16.0.19328.20178; Microsoft Corporation, Redmond, WA, USA), Clampfit version 10.4.2.0 (Molecular Devices, LLC, San Jose, CA, USA), custom-made Python analysis protocols, and GraphPad Prism 6.01 (GraphPad Software, LLC, Boston, MA, USA). Peak Na^+^ current densities were obtained by dividing the peak Na^+^ current amplitudes of each cell by the whole-cell membrane capacitance measured in the same cell. Na^+^ conductance at each test potential and in each cell was calculated using the following equation,

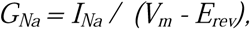

where *I_Na_* is the current amplitude at membrane voltage *V_m_* and *E_rev_* is the calculated Na^+^ reversal potential. Normalized conductances were then calculated as *G_Na_ / G_Na,Max_* where *G_Na,Max_* is the maximal conductance. Steady-state activation curves were derived by plotting the normalized conductance G_Na_ (G_Na_ / G_Na,Max_) as a function of the test potential and fitting the data using a Boltzmann equation of the following form:

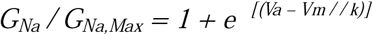

where *V_a_*is the membrane potential of half-maximal activation and *k* is a slope factor.

For steady-state inactivation, normalized current amplitudes *I_Na_ / I_Na,Max_* at each test potential in each cell were determined. Mean values were plotted as a function of the pre-pulse potential *V_m_*and fitted using a Boltzmann equation of the form:

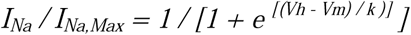

where *V_h_*is the voltage of half-maximal inactivation and *k* is a slope factor.

### Statistics

Normality of the data was assessed using the D’Agostino-Pearson test with a significance threshold of p < 0.05. N values throughout the text refer to the total number of cells analyzed in each experiment. As most of the data were not normally distributed, values are presented as median [first quartile, third quartile]. For current–action potential number relationships, data are shown as mean ± standard error of the mean and qPCR results are reported as mean ± standard deviation. For multiple comparisons, either non-parametric ANOVA (Kruskal– Wallis) with Dunn’s post-hoc tests or ordinary one-way ANOVA with Tukey’s post-hoc tests was performed, as appropriate. A two-factor ANOVA was used to evaluate the effect of XE-991 treatment across different cell lines and to assess potential interactions between genotype and drug application. qPCR analysis was performed on ΔΔCt values using a one-sample t-test against zero. All statistical analyses were conducted using GraphPad Prism 6.01 (GraphPad Software, LLC, Boston, MA, USA).

### Study approval

All experiments involving human induced pluripotent stem cells (hiPSCs) were approved by the Ethics Committee of the Montreal Neurological Institute, McGill University (IRB Number: MPE-CUSM-15-915 and ND02.045) and conducted in accordance with institutional and national ethical guidelines. Written informed consent was obtained from all donors of the original cell lines. The experiments were also in line with the vote of the Ethical Committee of the Medical Faculty of the University of Tübingen and the University Hospital Tübingen (428/2023BO2).

## Results

Human iPSC-derived neurons were generated from two SCA27B patients - one monoallelic patient, carrying one expanded repeat allele (391 plus a 19 GAA•TTC repeat allele), and one biallelic patient, carrying two expanded repeat alleles (290 and 315 GAA•TTC repeats) - as well as from two iPSC lines from healthy individuals (9/59 GAA•TTC repeats), using a Neurogenin 2 (NGN2) overexpression-based differentiation approach. *FGF14* mRNA levels and electrophysiological parameters were determined as described in the methods section.

### Clinical Case Vignettes

#### Case B6440 (mono-allelic)

Cell line B6440 was obtained from a 57-year-old male subject of French-Canadian descent. SCA27B started at age 40 years, with episodic ataxia. The episodes were characterized by gait ataxia, cerebellar dysarthria, and diplopia. Some episodes were triggered by alcohol intake and fatigue. The patient experienced about 1 episode per day, each lasting 2-3 hours. Treatment with acetazolamide did not provide benefit. At time of last assessment (age 63 years), he still reported no permanent symptoms or disability (SARA score 0/40 points). His clinical examination showed downbeat nystagmus and horizontal gaze-evoked nystagmus. Brain MRI was unremarkable.

#### Case B4796 (biallelic)

Cell line B4796 was obtained from a 68-year-old male subject of French-Canadian descent. His first symptoms started at age 52 years with episodes of gait ataxia, cerebellar dysarthria, and diplopia. He subsequently developed progressive gait ataxia at age 54 years, followed by appendicular ataxia and dysarthria. Acute episodes were triggered by alcohol intake, bright light, and fatigue. Acetazolamide provided no benefit. The patient started using a cane at age 55 and was wheelchair-bound by age 63 years. At time of last assessment at age 68, the neurological examination was remarkable for saccadic pursuit, horizontal gaze-evoked nystagmus, cerebellar dysarthria, upper and lower limb ataxia, and gait ataxia (SARA score 32.5/40; annualized SARA progression rate 2 points/year). The patient had hyperreflexia, but no spasticity. He had no clinical signs of polyneuropathy. Brain MRI at age 62 showed mild vermis atrophy (anterior lobe), no hyperintensity of the superior cerebellar peduncles, and a few scattered non-specific T2/FLAIR hyperintense lesions of the subcortical white matter. Due to the severity of his condition, the patient requested medical aid in dying at age 69.

### Reduced *FGF14* mRNA levels and neuronal excitability

The *FGF14* gene comprises five exons, with a deep intronic GAA•TTC repeat between the first and second exon of isoform 1b (Fig 1A). *FGF14-1b* mRNA levels were measured by quantitative polymerase chain reaction (qPCR) using SYBRgreen dye for double-strand detection. This analysis revealed a significant reduction in *FGF14-1b* mRNA levels of ∼40% in monoallelic (Fig 1B; mean ± SD *FGF14-1b* transcript expression of monoallelic relative to control, 0.58 ± 0.14, n = 4, p = 0.036, one-sample t-test against zero on ΔΔCt values) and ∼30% in biallelic patient-derived neurons compared to controls (mean ± SD *FGF14-1b* transcript expression of biallelic relative to control, 0.75 ± 0.11, n = 3, p = 0.046, one-sample t-test against zero on ΔΔCt values).

**Figure 1.**
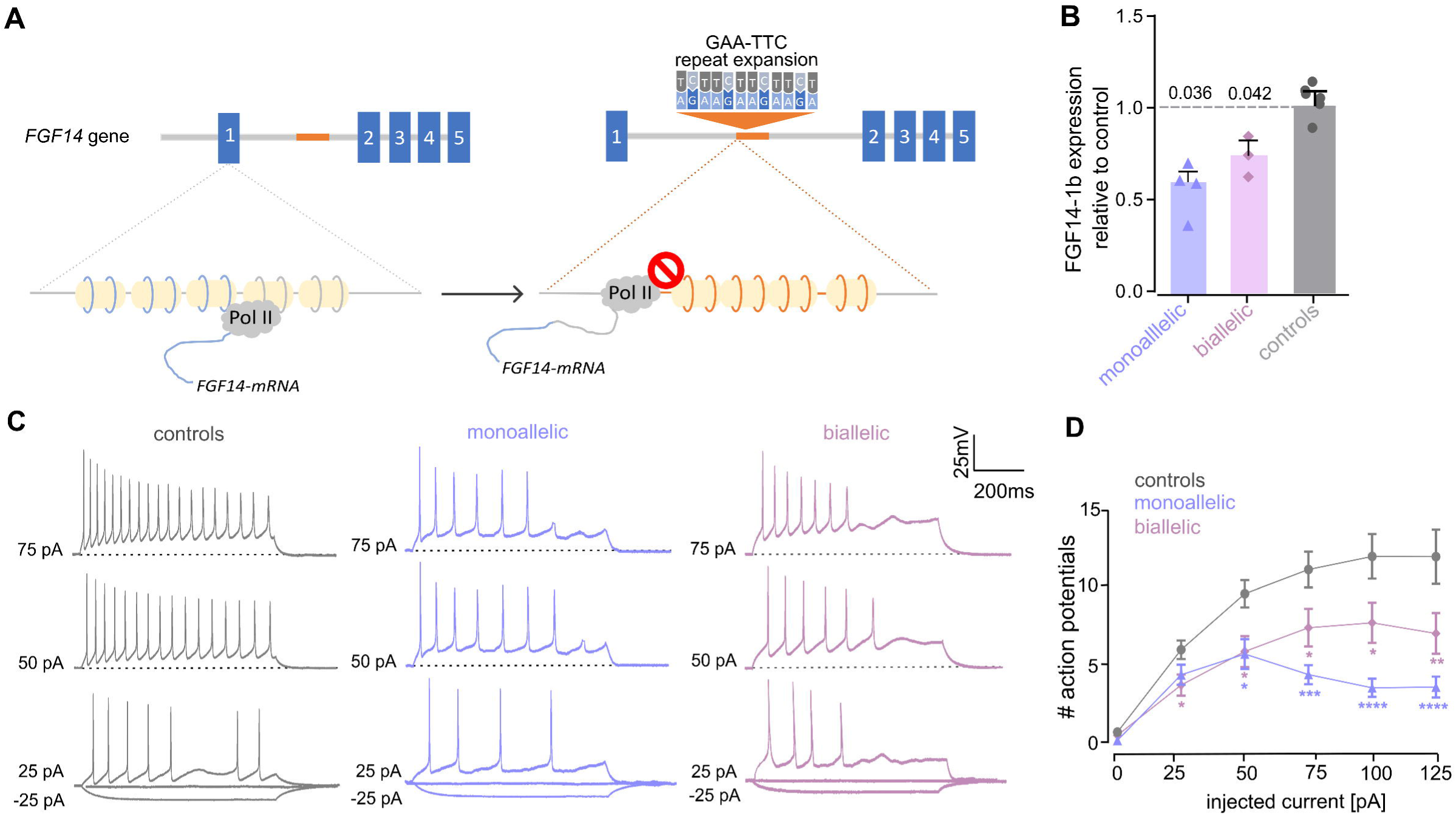
Reduced *FGF14-1b* transcription and reduced excitability in SCA27B patient-derived neurons. **(A)** Schematic representation of the *FGF14* gene with five exons (numbered blue boxes) and a deep intronic GAA•TTC repeat expansion located between the first and second exon. Below, orange circles indicate the GAA•TTC repeat sequence wrapped around histones (yellow ovals). RNA polymerase II is hindered to progress through the repeat expansion locus. **(B)** Expression of *FGF14-1b* mRNA by quantitative RT-PCR normalized to expression of *FGF14-1b* mRNA in controls in the monoallelic cell line (n = 4) and in the biallelic cell line (n = 3). Data are reported as mean ± SD and statistical analysis was performed on ΔΔCt values using a one-sample t-test against zero. **(C)** Representative whole-cell current clamp recordings of action potential series from NGN2-differentiated neurons, derived either from iPSC-lines from two normal controls (left; grey), from a SCA27B patient with a monoallelic (middle; blue) or a biallelic (right; pink) GAA•TTC repeat expansion in the *FGF14* gene upon injection of - 25, 0 and 25 pA (bottom), 50 pA (middle) and 75 pA (top). Dashed lines show holding of –70 mV. **(D)** Action potential counts are shown as a function of injected current for control neurons (pooled data), neurons carrying a monoallelic, or biallelic repeat expansion. Data are presented as mean ± SEM with sample sizes as follows: controls, n = 87; heterozygous, n = 44; biallelic, n = 27. Asterisks indicate adjusted p-values from a Kruskal–Wallis test followed by Dunn’s post hoc test (*p < 0.05, **p < 0.01, ***p < 0.001, ****p < 0.0001).

To assess excitability in patient-derived neurons, we recorded action potential (AP) firing by injecting stepwise depolarizing currents in 25 pA increments (Fig 1C). Both patient-derived neurons exhibited reduced AP firing, beginning at 25 pA in the monoallelic line and 50 pA in the biallelic line, compared to control neurons (Fig 1D). Moreover, we calculated the total number of APs across all current injections for each recorded neuron (designated as cumulative AP number), which was significantly reduced in both patient-derived lines relative to controls (exact values and statistics are provided in Table 1).

**Table 1.**
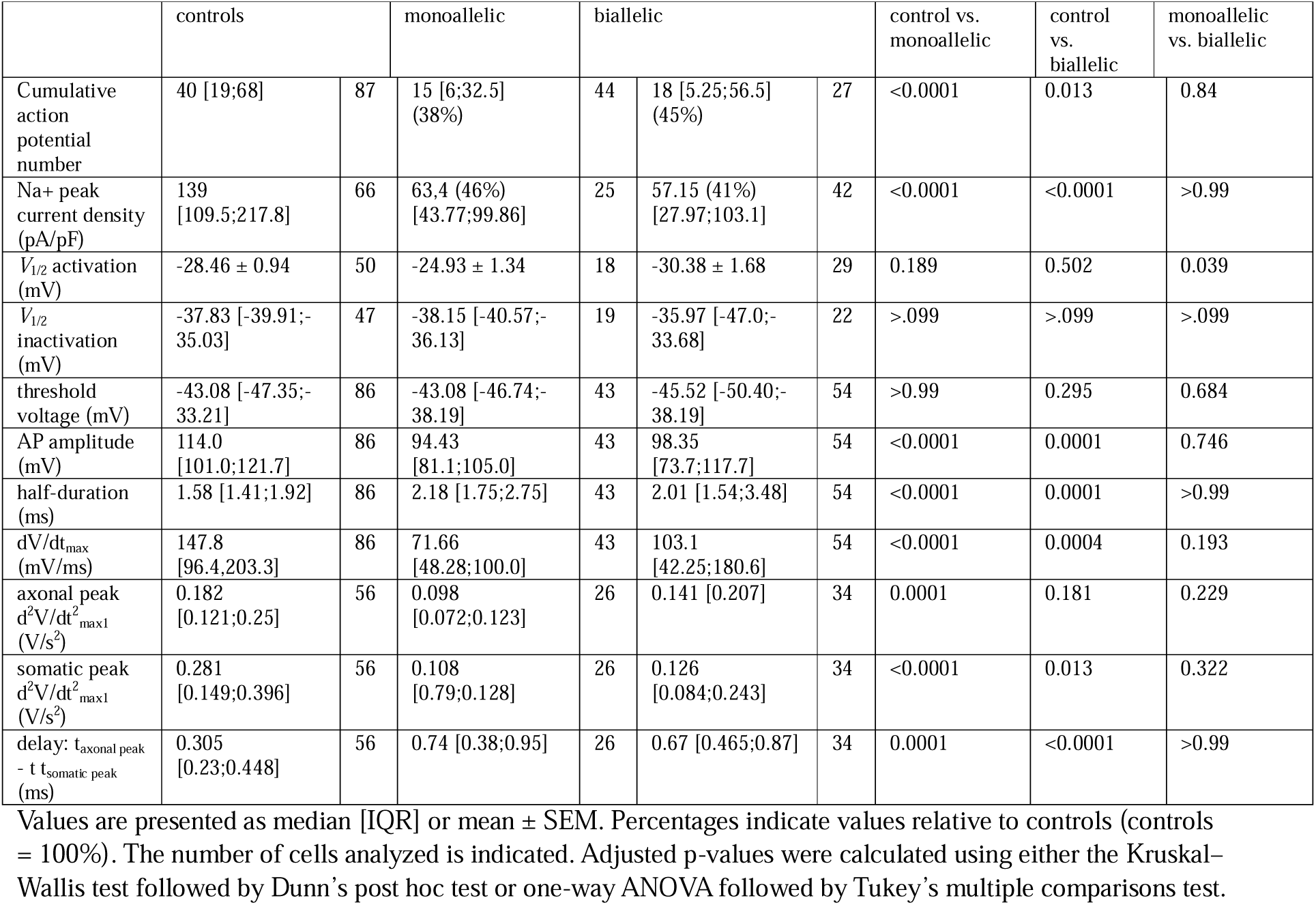
Electrophysiological parameters from spike count, Na^+^ currents and action potentials including first and second derivatives.

### Decreased Na^+^ current density

Previous studies have shown that FGF14 enhances Na^+^ current density at the axon initial segment (AIS) by promoting the clustering and localization of voltage-gated Na^+^ (Na_V_) channels through its binding to Na_V_ channels downstream of ankyrin-G [6, 17, 40]. FGF14 itself is subject to phosphorylation and serves as a target for multiple kinases, which modulate its activity and its regulation of Na_V_ channels [5, 41]. We therefore investigated whether the reduced neuronal excitability can be attributed to a reduction in the availability or function of Na_V_ channels. We therefore recorded NA^+^ currents under voltage-clamp conditions with increasing voltage steps from -60 mV to +20 mV in 10 mV increments, revealing a markedly reduced peak current density at -20 mV in both monoallelic and biallelic patient-derived neurons (Fig 2A, B and Table 1). Although peak Na^+^ current densities were reduced, voltage-dependent gating of the channels was not significantly affected (see inactivation and activation curves in Fig. 3C and V_1/2 activation_ and V 1/2 activation (mV) in Table 1).

**Figure 2.**
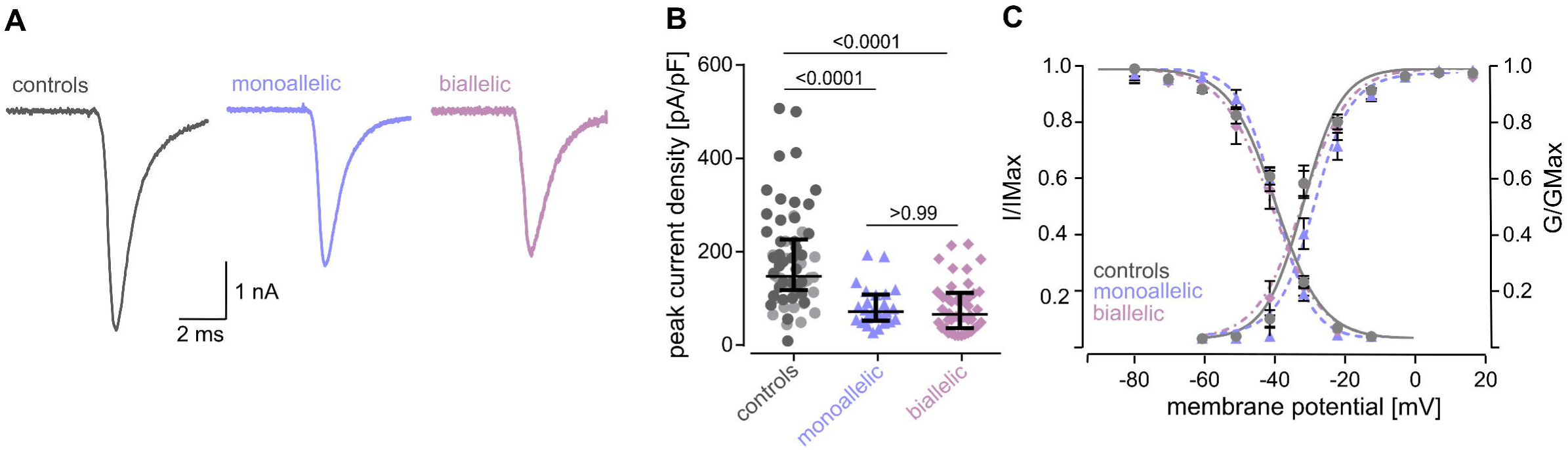
Decreased peak NA^+^ current density, but intact channel gating, in SCA27B patient-derived neurons. **(A)** Representative traces of NA^+^ currents recorded from controls (left; grey), monoallelic (middle; blue) and biallelic (right; pink) patient-derived neurons were elicited by voltage steps from –60 to 30 mV in 10 mV increments (potential –70 mV). **(B)** Scatter plots of the peak NA^+^ current density of neurons recorded at -20 mV in control, monoallelic and biallelic patient-derived neurons. Peak current density was significantly decreased for both monoallelic and biallelic without difference between patient lines. Data are presented as median [IQR] with sample sizes as follows: controls, n = 66; monoallelic, n = 25 and biallelic, n = 42. Adjusted p-values (Kruskal–Wallis test with Dunn’s post hoc correction) are shown in the figure. **(C)** Voltage-dependence of steady-state activation and inactivation show no change between control and patient cell lines. Lines represent Boltzmann functions fit to the data points, representing means ± SEM. Colour code as in (A).

**Figure 3.**
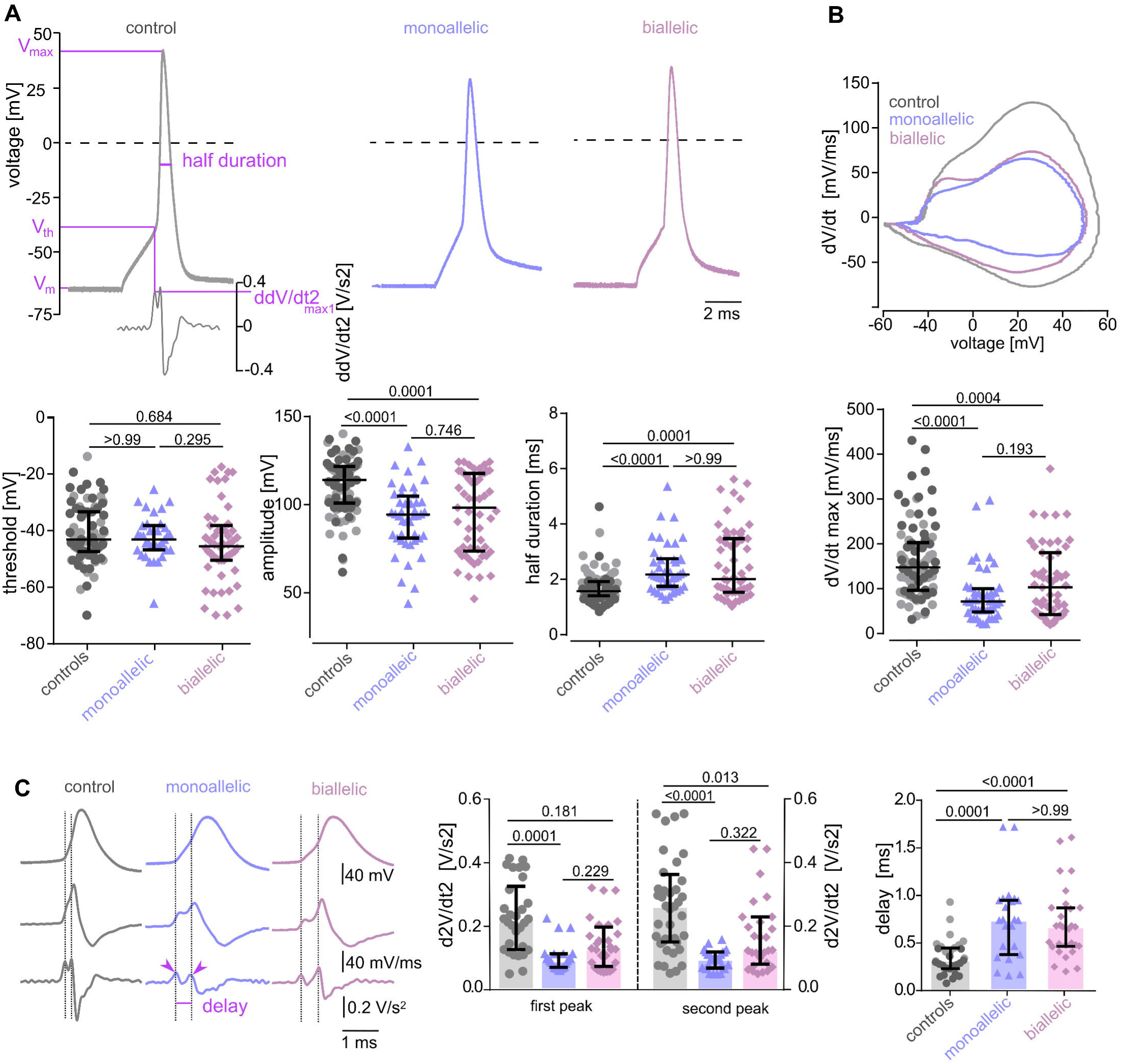
Slowed depolarization and delayed spike initiation in SCA27B patient-derived neurons. **(A)** *Top:* Example traces showing the parameters used to describe an action potential, including threshold voltage V_th_, amplitude (difference from V_max_ and V_m_) and half-duration. *Bottom*: Scatter plots of those parameters indicating no change in threshold voltage V_th_ but decreased amplitude and increased half-duration for both patient lines (controls: n = 86; monoallelic: n = 43 and biallelic: n = 54). **(B)** *Top:* Representative phase plots of dV/dt vs *V* for spikes generated from recordings as in (A). *Bottom:* Scatter plots of dV/dt_max_ showing decreased maximum upstroke velocity for both patient cell lines (controls: n = 86; monoallelic: n = 43 and biallelic: n = 54). **(C)** *Left:* Representative traces of spikes and their corresponding first (middle trace) and second (bottom trace) derivations of controls, monoallelic and biallelic patient-derived neurons generated from recordings as in (A). Black dotted lines illustrate the delay between the first (axonal) and the second (somatic) peak. *Middle:* Scatter plots of d^2^V/dt^2^_max1;2_ showing decreased axonal (first) and somatic (second) peak in the second derivation in monoallelic patient-derived neurons and decreased somatic peak in biallelic-patient derived neurons. *Right:* Delay between the axonal and somatic peak indicating attenuated spike propagation in both patient cell lines (analyzed from recordings, as in (A) controls: n = 56; monoallelic n = 26 and biallelic: n = 34). Data are presented as median [IQR]. Adjusted p-values (Kruskal–Wallis test followed by Dunn’s multiple comparisons test) are indicated in the figure.

### Delayed AP initiation and decreased membrane potential acceleration

To further elucidate Nav channel pathology in neurons derived from SCA27B patients and to investigate whether channels are lost or redistributed at the AIS, we analyzed AP parameters in more detail (Fig. 3, Table 1). Consistent with unaltered channel activation properties, the AP threshold (V□□) remained unchanged. In line with the observed reduction in peak NA^+^ currents, AP amplitudes were reduced in both monoallelic and biallelic patient-derived neurons (Fig 3A and Table 1). Phase plot analysis of the first derivative (dV/dt) revealed that patient-derived neurons also exhibited a decrease in maximal depolarization rate (dV/dt max, Fig 3B and Table 1), reflecting diminished inward NA^+^ current during the AP upstroke. As a consequence of slower and reduced regenerative depolarization, AP half-duration was prolonged, further supporting decreased NA^+^ current driving the upstroke (Fig 3A and Table 1).

Analysis of the second derivative of the AP (d²V/dt²) allowed distinction between axonally initiated and somatic spike components (Fig 3C). Two peaks were identified: the first peak, corresponding to the axonal spike (AIS initiation), and the second peak, representing the somatic spike [42, 43]. In the monoallelic line, the first axonal peak was selectively reduced, and the somatic peak was reduced in both patient lines, reflecting decrease in membrane potential acceleration. Moreover, both patient lines showed an increased latency between axonal and somatic peaks. This delayed AP initiation reflects either slower propagation time from the initiation at the AIS to the invasion at the soma or a shift in the in the spike initiation site [43].

Together, these data indicate that SCA27B neurons display a cascade of NA^+^ channel-dependent functional deficits, affecting both AP upstroke and axonal initiation, ultimately leading to reduced neuronal excitability. These deficits likely result from loss of functionally available NA^+^ channels at the AIS, without altering intrinsic channel gating properties.

### M-currents are not affected

In addition to its interaction surface with Nav channels, FGF14 has been shown to bind to KCNQ2 (K_V_7.2) channels [23]. Kv7.2 and Kv7.3 channels generate the M-current, a slowly activating and non-inactivating potassium current that plays a key role the control of neuronal excitability. We pharmacologically isolated the M-current by blockage of voltage-gated Na^+^ and Ca^2+^ channels and K^+^ channel K_V_1 and to K_V_3 families, using tetrodotoxin, Cd^2+,^ and 4-aminopyridine, and by using XE-991, a K_V_7 selective inhibitor. We then determined the XE-991-sensitive current by subtracting the XE-991-resistant component from the baseline current. XE-991 significantly decreased the remaining K^+^ current amplitudes, but there were no significant differences in the basal and XE-991-resistant K^+^ current amplitudes between control, monoallelic, and biallelic patient-derived neurons and no significant differences in the XE-991 sensitive component (Fig 4 and Table 2). This suggests that, at least in this model system, K_V_7.2 channels are not affected in impaired excitability.

**Figure 4.**
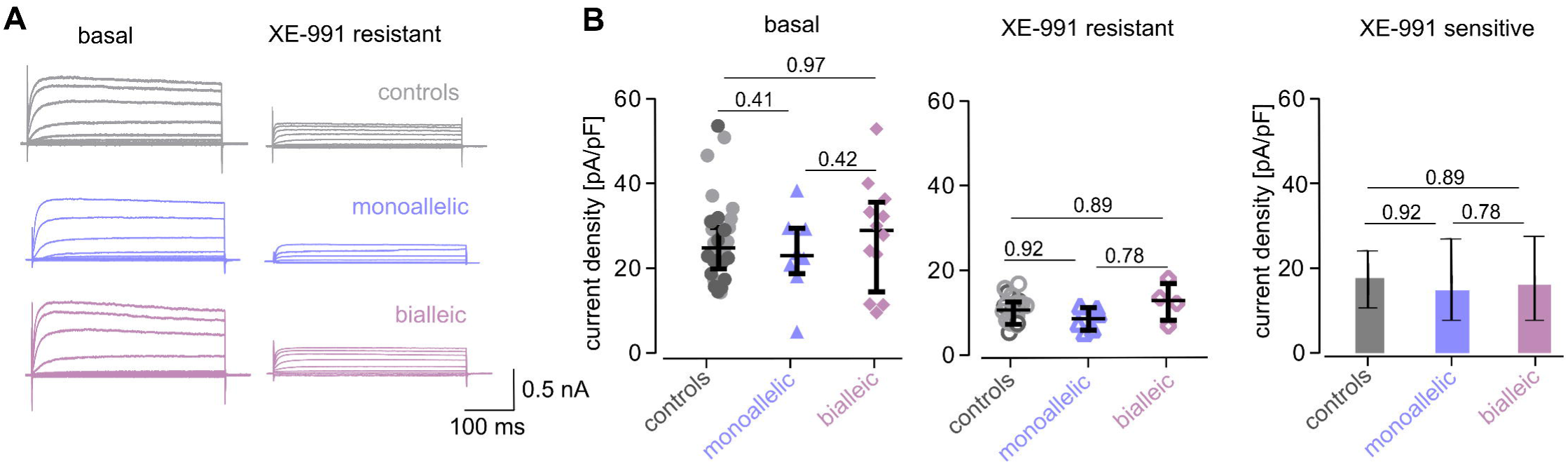
XE-991-sensitive KCNQ outward currents are not impaired in SCA27B patient-derived neurons. **(A)** Exemplary K+ currents from controls (top; grey), monoallelic (middle; blue) and biallelic (bottom; pink) before (left) and after (right) application of the KCNQ-channel blocker XE-991. **(B)** Left: Scatter plots of basal and XE991-resistant K+ current amplitudes from the indicated cell lines represented as median ± IQR. Right: XE-991 sensitive K+ current amplitudes showed no significant differences between cell lines. Data are represented as mean +95% confidence interval. controls: n = 37, monoallelic: n = 8, biallelic: n = 13 and with XE-991: controls: n = 19, heterozygous: n = 6, biallelic: n = 4; two-way ANOVA: treatment effect p < 0.0001, cell line effect p = 0.41 and interaction p = 0.86.

**Table 2.**
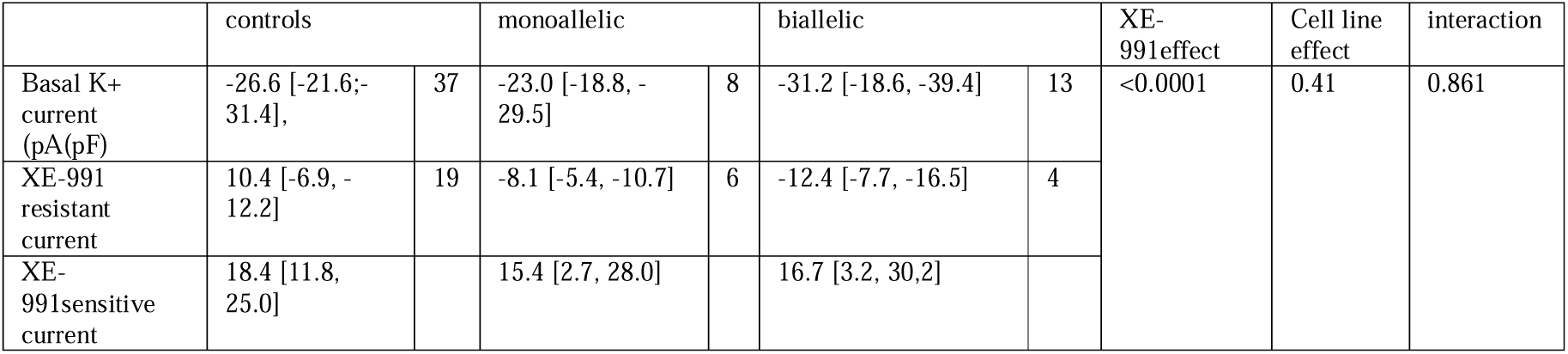
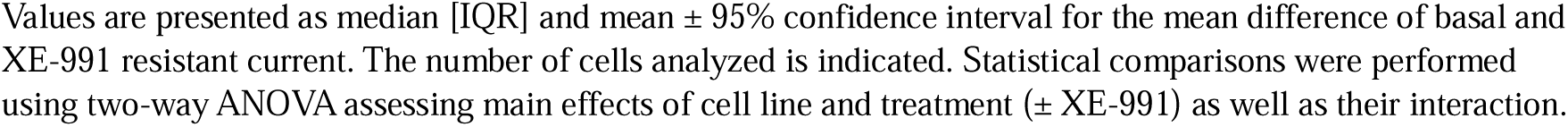
KCNQ currents are not affected.

|  | controls |  | monoallelic |  | biallelic |  | XE-991effect | Cell line effect | interaction |
| --- | --- | --- | --- | --- | --- | --- | --- | --- | --- |
| Basal K<sup>+</sup> current (pA/pF) | -26.6 [-21.6;-31.4], | 37 | -23.0 [-18.8, -29.5] | 8 | -31.2 [-18.6, -39.4] | 13 | <0.0001 | 0.41 | 0.861 |
| XE-991 resistant current | 10.4 [-6.9, -12.2] | 19 | -8.1 [-5.4, -10.7] | 6 | -12.4 [-7.7, -16.5] | 4 |  |  |  |
| XE-991sensitive current | 18.4 [11.8, 25.0] |  | 15.4 [2.7, 28.0] |  | 16.7 [3.2, 30.2] |  |  |  |  |

Values are presented as median [IQR] and mean ± 95% confidence interval for the mean difference of basal and XE-991 resistant current. The number of cells analyzed is indicated. Statistical comparisons were performed using two-way ANOVA assessing main effects of cell line and treatment ( ± XE-991) as well as their interaction.

### Reduced *FGF14* mRNA levels can be restored by Syn-TEF1

As we hypothesized that transcriptional deficiency is the key upstream event of SCA27B pathophysiology, we next targeted transcription failure by the small molecule synthetic transcription elongation factor 1 (Syn-TEF1). We expected that restoration of transcriptional deficiency would consequently also restore the “downstream events” of SCA27B pathophysiology, i.e. impaired neuronal excitability and NA^+^current density. Administration of Syn-TEF1 revealed a significant upregulation in *FGF14-1b* mRNA levels of ∼70% in monoallelic and ∼50% in biallelic patient-derived neurons compared to untreated neurons (Fig 6C; mean ± SD *FGF14-1b* transcript expression in monoallelic and biallelic patient-derived neurons upon Syn-TEF1 treatment relative to untreated condition, 1.73 ± 0.36, n = 4, p = 0.017 and 1.47 ±0.32, n = 3, p = 0.046, one-sample t-test against zero on ΔΔCt values). In contrast, Syn-TEF1 treatment had no effect on *FGF14-1b* mRNA levels in neurons derived from healthy controls (mean ± SD *FGF14-1b* transcript expression of controls upon syn-TEF1 treatment relative to untreated controls: 1.02 ± 0.06, n = 2, p = 0.705), suggesting that it does not lead to (potentially dysfunctional/toxic) “overexpression” of *FGF14*, which would potentially limit its therapeutic application.

**Figure 5.**
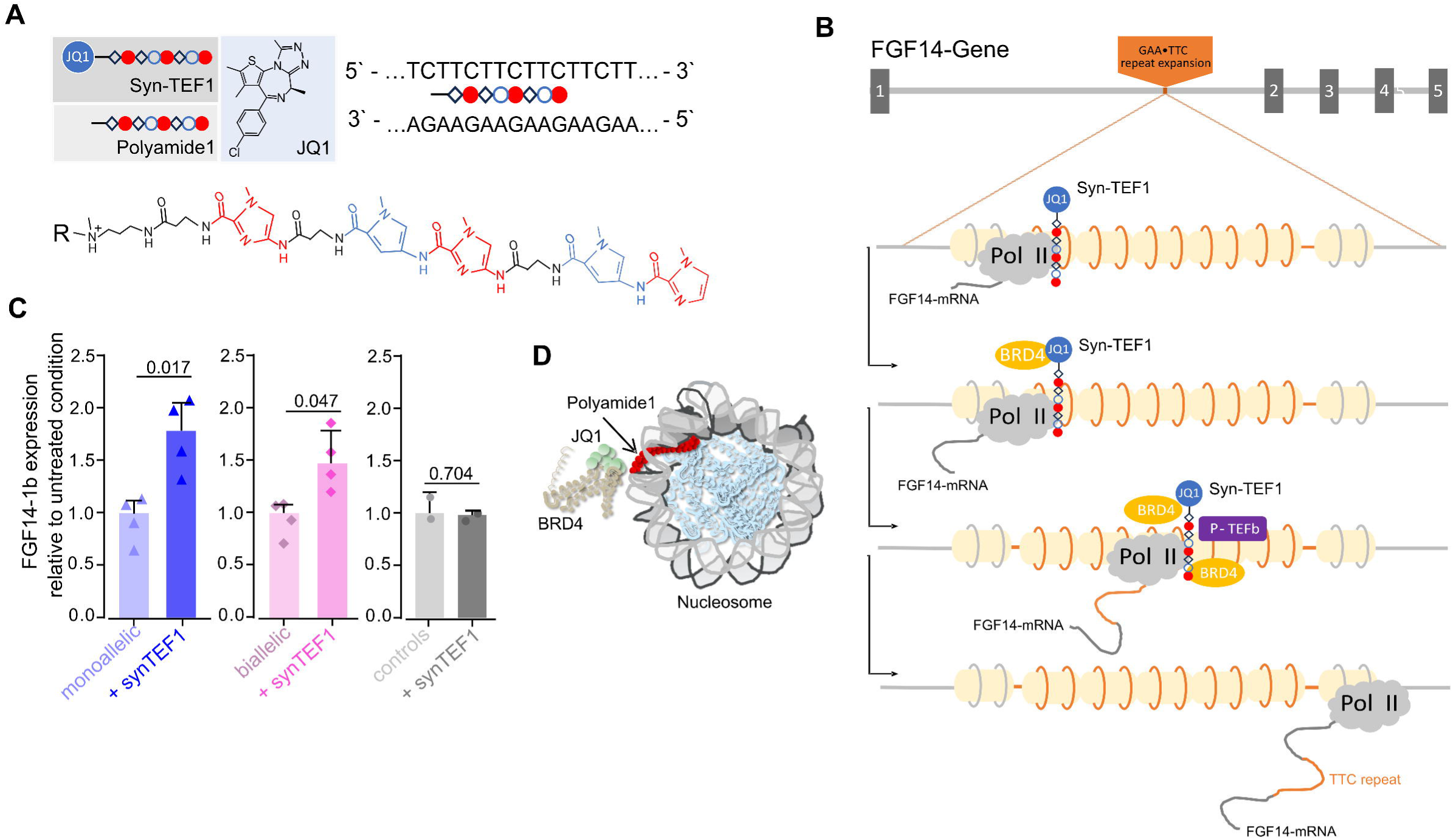
Synthetic transcription elongation factor 1 (SynTEF1) restores FGF14 mRNA expression in SCA27B patient-derived neurons. **(A)** Top left: Schematic representation of Syn-TEF1, consisting of a polyamide (colored chain) connected to the small molecule JQ1 (blue circle, bromodomain inhibitor) with a synthetic linker (. The molecular structure of its main components - the polyamide part and JQ1 – are presented also in more detail in isolation. Top right: binding mode of the polyamide to G-C and A-T base pairs, respectively. Bottom: Chemical structure of the polyamide consisting of DNA-specific sequence of N-methylimidazole (red) and N-methylpyrrole (blue), which are linked via β-alanine (black). **(B)** Model of the assumed cascade of interactions and molecular processes initiated by synthetic transcription elongation factor 1 (Syn-TEF1). Fibroblast Growth Factor (*FGF14)* gene with five exons (grey squares) and a GAA·TTC repeat expansion in the first intron, (orange) which constitutes a barrier to RNA polymerase II (Pol II, cloud-shapes). Syn-TEF1 binds at the repeat expansion locus (orange circles) via its polyamide 1 component (pearl line) and, through JQ1 (blue circle), recruits the transcriptional regulators bromodomain-containing protein 4 (BRD4; orange oval) and positive transcription elongation factor b (P-TEFb; purple square). This recruitment enhances Pol II elongation through the repressive chromatin structure, eventually allowing successful completion of *FGF14* transcription (adapted from [1]). **(C)** Relative expression of *FGF14-1b* mRNA in patient-derived neurons upon treatment with 1 µM syn-TEF1 by quantitative RT-PCR normalized to expression of *FGF14-1b* mRNA in untreated neurons. Left: monoallelic cell line (n = 4), middle: biallelic cell line (n = 3), right: control (n = 2). Data are reported as mean ± SD and statistical analysis was performed on ΔΔCt values using a one-sample t-test against zero. **(D)** The polyamide (red pearl) hybridises to nucleosomal DNA (symbolized by grey spirals), and JQ1 recruits BRD4 (Adapted from [1]).

**Fig. 6.**
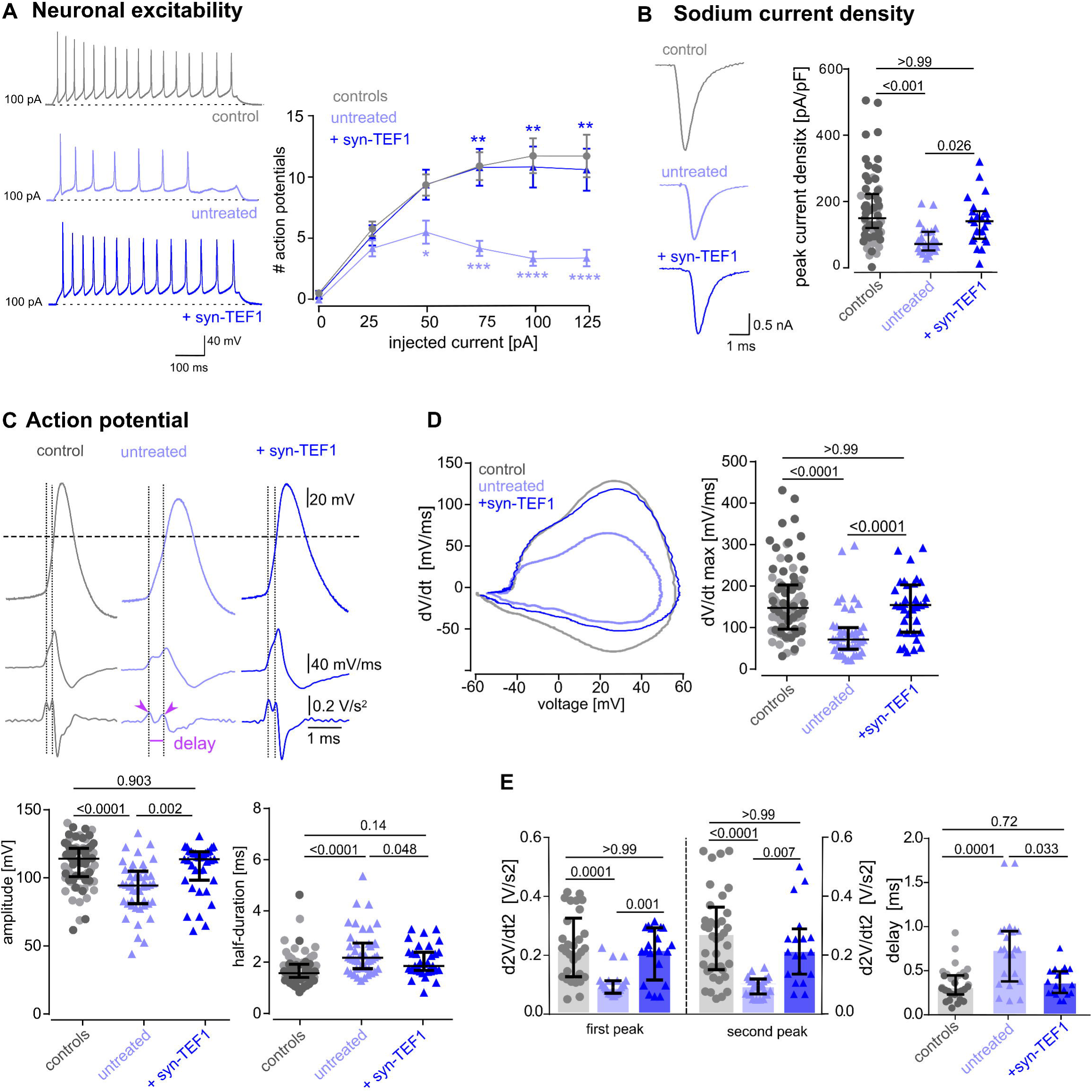
Syn-TEF1 restores excitability, NA^+^ current density and AP initiation in monoallelic patient-derived neurons. **(A)** *Left:* Representative whole-cell current clamp recordings of action potential series from control (top; grey), from untreated (middle; light blue) or syn-TEF1 treated (bottom; dark blue) monoallelic patient-derived neurons. Dashed lines show holding of –70 mV. *Right:* Action potential (AP) counts are shown as a function of injected current showing significantly increased AP firing upon syn-TEF1 treatment. Data are presented as mean ± SEM with sample sizes as follows: controls, n = 87; untreated, n = 44; treated, n = 38. Light blue asterisks: controls vs untreated condition; dark blue asterisks: untreated vs treated condition. **(B)** *Left:* Representative traces of NA^+^ currents recorded at -20 mV from the indicated cell lines and conditions. *Right:* Scatter plots of the peak NA^+^ current density recorded at -20 mV which was restored upon syn-TEF1 treatment. Data are presented as median ± IQR (controls, n = 66; untreated, n = 25 and +syn-TEF1terated, n = 24**). (C)** *Top:* Representative traces of spikes and their corresponding first (middle trace) and second (bottom trace) derivations. Vertical dotted lines illustrate the delay between the first (axonal) and the second (somatic) peak. Horizontal dotted line indicates potential at 0 mV. *Bottom:* Scatter plots of AP amplitude and half-duration which both are restored upon syn-TEF1 treatment (controls: n = 86; untreated: n = 43 and +syn-TEF1treated: n = 35). **(D)** *Left:* Representative phase plots of dV/dt vs *V* for spikes generated from recordings as in (C). *Rigth:* Scatter plots of dV/dt_max_ showing restored maximum upstroke velocity upon syn-TEF1 treatment (controls: n = 66; untreated: n = 43 and +syn-TEF1terated: n = 35). **(E)** *Left:* Scatter plots of d^2^V/dt^2^_max1;2_ showing restored axonal (first) and somatic (second) peak in the second derivation *Right:* Delay between the axonal and somatic peak was restored upon syn-TEF1 treatment (analyzed from recordings, as in (C) controls: n = 56; untreated: n = 26 and syn-TEF1 treated: n = 24). Data are presented as median [IQR]. Values from controls are identical as shown in Fig. 2,3 and 5. Adjusted p-values (Kruskal–Wallis test followed by Dunn’s multiple comparisons test) are indicated in the figure (*p < 0.05, **p < 0.01, ***p < 0.001, ****p < 0.0001).

### Neuronal excitability, NA^+^ current density and slowed AP initiation are restored by Syn-TEF1

To test the functional consequences of Syn-TEF1-driven upregulation of *FGF14-1b* expression, we recorded trains of AP by increasing current injections upon at least three days treatment with 1 µM Syn-TEF1. We observed a significant upregulation of AP firing and the cumulative number of APs upon treatment both for the monoallelic (Fig 7 and Table 3) and the biallelic patient-derived neurons (Fig 8 and Table 4) with restoration to similar levels as observed in control neurons.

**Fig. 7.**
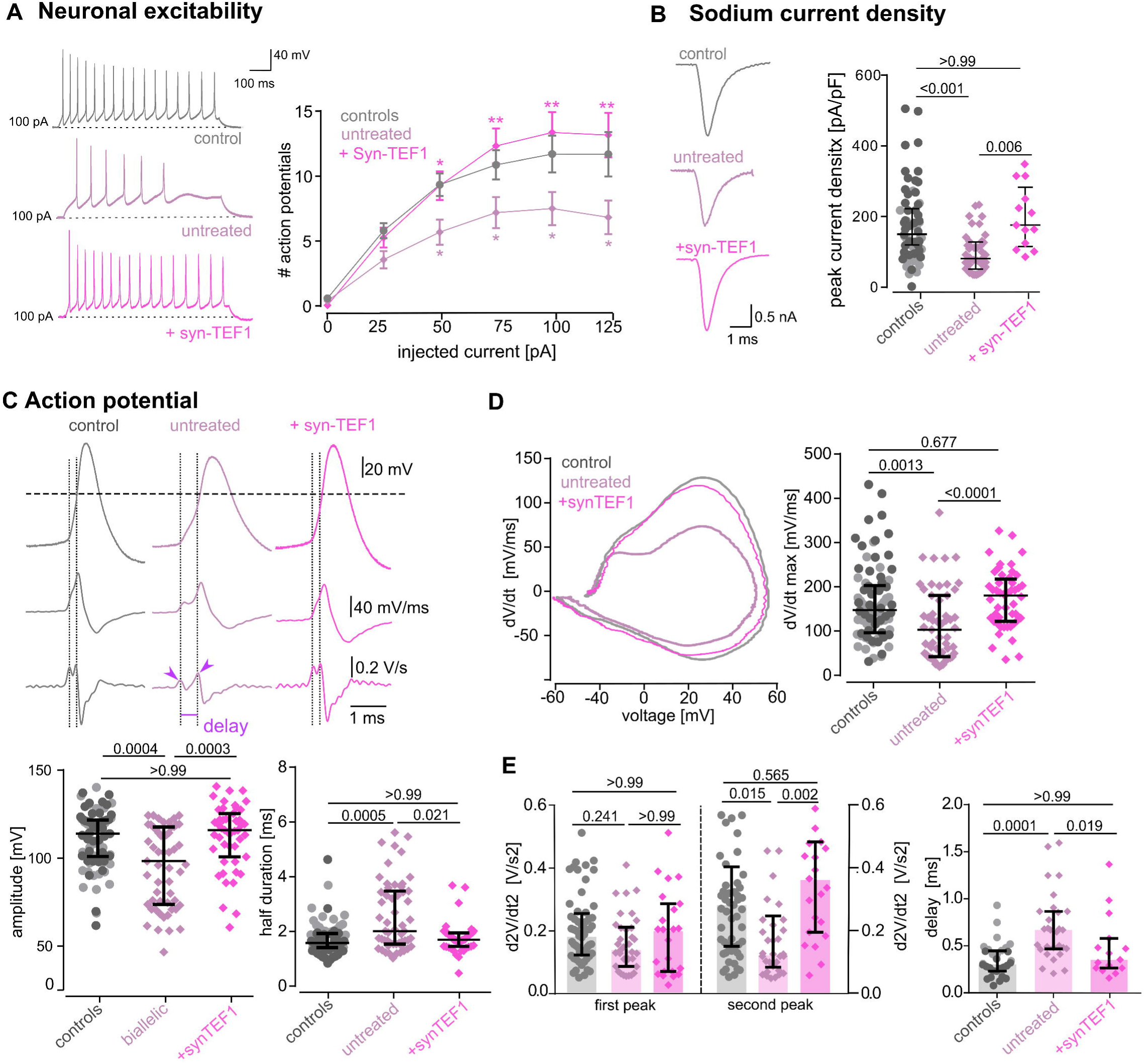
Syn-TEF1 restores excitability, NA^+^ current density and AP initiation in biallelic patient-derived neurons. **(A)** *Left:* Representative whole-cell current clamp recordings of action potential series from control (top; grey), from untreated (middle; pink) or syn-TEF1 treated (bottom; neon pink) biallelic patient-derived neurons. Dashed lines show holding of –70 mV. *Right:* Action potential (AP) counts are shown as a function of injected current showing significantly increased AP firing upon syn-TEF1 treatment. Data are presented as mean ± SEM with sample sizes as follows: controls, n = 87; untreated, n = 27; treated, n = 37. Pink asterisks: controls vs untreated condition; neon pink asterisks: untreated vs treated condition. **(B)** *Left:* Representative traces of NA^+^currents recorded at -20 mV. *Right:* Scatter plots of the peak NA^+^ current density recorded at -20 mV which was restored upon syn-TEF1 treatment. Data are presented as median ± IQR (controls, n = 66; untreated, n = 42 and +syn-TEF1terated, n = 16**). (C)** *Top:* Representative traces of spikes and their corresponding first (middle trace) and second (bottom trace) derivations. Vertical dotted lines illustrate the delay between the first (axonal) and the second (somatic) peak. Horizontal dotted line indicates potential at 0 mV. *Bottom:* Scatter plots of AP amplitude and half-duration which both are restored upon syn-TEF1 treatment (controls: n = 86; untreated: n = 54 and +syn-TEF1 treated: n = 47). **(D)** *Left:* Representative phase plots of dV/dt vs *V* for spikes generated from recordings as in (C). *Rigth:* Scatter plots of dV/dt_max_ showing restored maximum upstroke velocity upon syn-TEF1 treatment (controls: n = 86; untreated: n = 54 and +syn-TEF1terated: n = 47). **(E)** *Left:* Scatter plots of d^2^V/dt^2^_max1;2_ showing restored somatic (second) peak in the second derivation *Right:* Delay between the axonal and somatic peak was restored upon syn-TEF1 treatment (analyzed from recordings, as in (C) controls: n = 56; untreated: n = 34 and syn-TEF1 treated: n = 23). Data are presented as median [IQR]. Values from controls are identical as shown in Fig. 2,3 and 5. Adjusted p-values (Kruskal–Wallis test followed by Dunn’s multiple comparisons test) are indicated in the figure (*p < 0.05, **p < 0.01, ***p < 0.001, ****p < 0.0001).

**Table 3.**
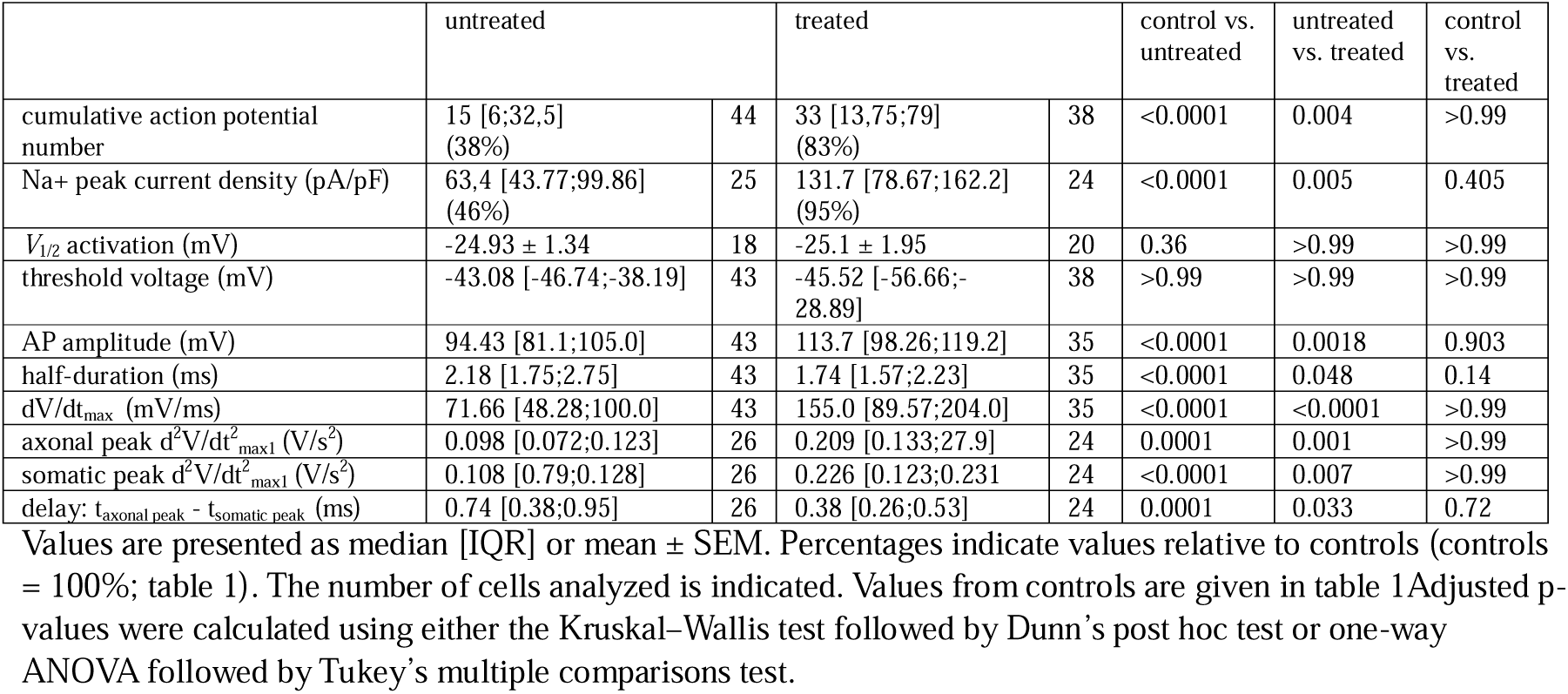
syn-TEF1 effect in monoallelic patient-derived neurons.

**Table 4.**
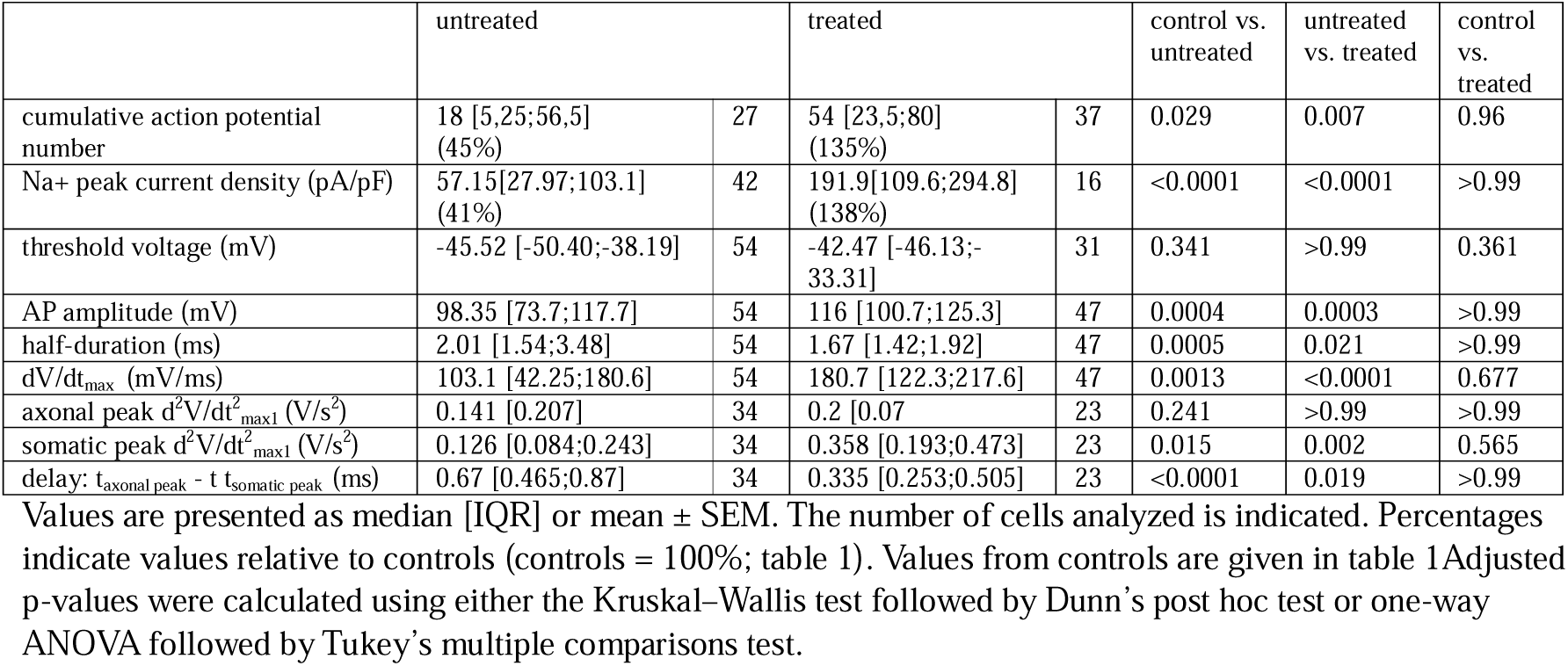
syn-TEF1 effect in biallelic patient-derived neurons.

Analyses of peak Na^+^ currents upon Syn-TEF1 treatment showed significantly increased amplitudes of Na^+^ peak current density for both monoallelic (Fig 7 and Table 3) and biallelic (Fig 8 and Table 4) patient-derived neurons, with restoration to similar levels as observed under control conditions. Importantly, the voltage-dependence of activation and inactivation, which did not differ between patient-derived and control neurons at baseline (Table 1), remained unaltered upon Syn-TEF1 treatment (Tables 3 and 4). Similarly, the action potential threshold was unchanged by treatment (Tables 3 and 4). These observations strongly indicate that Syn-TEF1 selectively restores the deficits in neuronal excitability and AP dynamics without inducing a pathological increase of FGF14 expression or an over-compensation of functional parameters. Following the restoration of peak Na^+^ current density and AP firing frequency, we next evaluated whether Syn-TEF1 treatment could rescue additional action potential properties. Detailed analysis revealed that AP amplitude and half-duration, as well as the maximal depolarization rate (V□/dt□□□), were restored in patient-derived neurons (Fig 7C, D and Fig 8C, D). In both monoallelic and biallelic neurons, Syn-TEF1 treatment normalized the previously observed delay between axonal and somatic action potential peaks, effectively resolving the slowed AP initiation. Moreover, the reduced acceleration of voltage change at the axonal peak, which was particularly pronounced in monoallelic patient-derived neurons and the somatic peak amplitude of the second derivative, previously reduced in both patient lines, was similarly rescued.

## Discussion

Our study demonstrates that transcriptional deficiency of FGF14, caused by GAA•TTC repeat expansions, represents the key upstream event driving impaired neuronal excitability in SCA27B patient-derived neurons. By selectively targeting this transcriptional block with Syn-TEF1, we achieved robust restoration of Na^+^ current density, slowed AP initiation and neuronal excitability, effectively rescuing the downstream functional deficits of the SCA27B pathophysiological cascade. These findings provide evidence that reduced FGF14 transcription is one principal driver of neuronal dysfunction in SCA27B and establish a proof-of-concept for a gene-targeted, disease-modifying therapeutic approach in SCA27B

### Transcriptional deficiency of *FGF14* as an upstream event of SCA27B pathophysiology

We observed downregulation of the *FGF14-1b* isoform in NGN2-induced human neurons derived from both monoallelic and biallelic SCA27B patient lines, suggesting that the GAA•TTC repeat expansion directly impairs transcription. This finding corroborates and extends recent reports demonstrating reduced *FGF14-1b* mRNA and protein expression in both postmortem cerebellar tissue and patient-derived motor neurons [1]. Notably in our model no differences in expression were observed between the biallelic patient line - carrying two expanded alleles - and the monoallelic patient line harbouring an expansion that was approximately ∼90 repeats longer than the largest expanded allele in the biallelic cell line.

In Friedreich’s ataxia (FA) expanded intronic GAA•TTC repeat tracts, often exceeding those reported in SCA27B, have been shown to induce the formation of triple-helical DNA (H-DNA) structures and R-loops, which hinder RNA polymerase II elongation, leading to transcriptional repression [44–47]. These structures often co-occur with epigenetic alterations, including heterochromatin formation and histone modifications, contributing to partial gene silencing [48, 49]. A similar GAA•TTC repeat driven transcriptional repression mechanism might take place in SCA27B yet warrants further specific mechanistic confirmation.

### Reduced Na^+^ currents and excitability in SCA27B patient iPSC-derived neurons

In line with the essential role of FGF14 in neuronal excitability, we observed that the transcriptional deficiency of FGF14 is associated with a marked reduction in intrinsic excitability in patient-derived neurons. These results provide the first functional evidence in human neurons, directly supporting and extending previous findings in mouse neurons, where knock-down of intracellular FGF14 similarly impaired neuronal excitability[10, 13], suggesting that this electrophysiological cellular phenotype is conserved across different cell types and species.

To understand the cellular basis of this impaired excitability, we next investigated voltage-dependent NA^+^and potassium currents carried by those channels that had been shown before to interact with FGF14 [23, 40]. Whole-cell patch-clamp recordings revealed a significant reduction in peak Na^+^ current density in patient-derived neurons, while the voltage dependence of activation and inactivation was not altered. While the unchanged AP threshold further supports preserved channel gating parameters, the observed reduction in AP amplitude and maximal depolarization rate (dV/dt max) points to a diminished availability of functional NA^+^ channels. This interpretation aligns with previous studies showing that FGF14 knockout or knockdown indeed leads to decreased NA^+^ channel density and disrupted distribution at the AIS [7, 13]. Yet, it contrasts with additional reports suggesting also altered gating properties in FGF14-deficient models [7, 15, 17]. More detailed analysis of the AP revealed a reduction of the second-derivative peaks, suggesting decreased membrane potential acceleration in patient neurons. Moreover, increased latency between the axonal and the somatic second derivative peaks points to delayed AP initiation and somatic invasion of the AP. These findings align with previous studies demonstrating that AP initiation critically depends on the high density and proper organization of Na_V_ channels at the AIS. Disruption of this compartment, even without major changes in channel gating, can lead to reduced membrane potential acceleration and delayed AP initiation [42, 43].

In addition to its role in NA^+^ channel positioning and function, FGF14 has also been shown to interact directly with voltage-gated potassium channels, particularly of the K_V_7 (KCNQ) family generating the M-current, which is critical for determining the subthreshold membrane potential and neuronal firing rate, membrane repolarization and action potential recovery [23, 50, 51]. However, when assessing M-currents in our human iPSC-derived neurons, no significant differences between lines from SCA27B patients and controls were observed, suggesting that M-current function is preserved in this model.

Our findings together with previous results thus suggest that FGF14 haploinsufficiency disrupts the functional availability of Nav channels. This leads to decreased Na^+^ current density and consequently reduced depolarization rates culminating in reduced neuronal excitability and firing frequency.

### Syn-TEF1 restores transcription deficiency and, consequently, Na^+^ currents and neuronal excitability

Synthetic transcription elongation factors (Syn-TEFs) are engineered fusion proteins that promote productive transcription through regions of expanded nucleotide repeats by counteracting repeat-mediated transcriptional stalling [26, 34]. They are composed of a sequence-specific DNA-binding polyamide, which is covalently linked to the small molecule JQ1 via a flexible PEG spacer through an amide bond. The BET-bromodomain-inhibitor JQ1 engages the transcription elongation machinery, allowing reactivation of genes silenced by expanded repeats (Fig 7). Originally tested in Friedreich’s ataxia, in which GAA•TTC repeat expansions (600-1100 GAA•TTC repeats) in the first intron of the frataxin gene impair transcription, Syn-TEF1 has been shown to restore frataxin mRNA levels and improve disease-relevant outcomes in cellular models [26, 52]. Given that SCA27B is likewise caused by an intronic GAA•TTC repeat expansion (250-500 GAA•TTC repeats) - here in the *FGF14* gene, with TTC as the sense strand (unlike in FA) - we hypothesized that Syn-TEF1 might also alleviate pathogenic transcriptional repression in this context [53]. The Syn-TEF1 derivative DT-216P2 is now tested in a Phase□1/2 clinical trial (ClinicalTrials.gov Identifier: NCT06874010), thus highlighting the translational clinical potential of this transcriptional elongation–based strategy for SCA27B.

Our data show that Syn-TEF1 treatment of SCA27B patient-derived neurons leads to a significant upregulation of the disease-relevant *FGF14*-transcript isoform 1b, restoring its expression towards similar levels as seen in controls. This observation corroborates the view that the reduction in *FGF14* mRNA levels is indeed due to repeat-mediated transcriptional repression [1]. Moreover, the absence of upregulation of *FGF14* mRNA in control cell lines suggests that Syn-TEF1 selectively targets loci with stalled RNA polymerase (rather than broadly activating all GAA•TTC repeat–containing sequences), again consistent with a requirement for transcriptional pausing as its mechanism of action. RNA-Seq analyses in FA further show that Syn-TEF1 exhibits minimal off-target effects, with little to no upregulation of other genes containing GAA repeats [27

The molecular restoration of *FGF14* transcription by Syn-TEF1 was accompanied by a rescue of key functional deficits previously observed in SCA27B patient-derived neurons. Treatment led to normalization of action potential firing frequency and peak NA^+^ current density in both monoallelic and biallelic neurons, without altering the voltage-dependence of activation or inactivation, or AP threshold. Importantly, additional properties of neuronal excitability, including AP amplitude, half-duration, and maximal depolarization rate, were similarly restored, and the previously observed delay between axonal and somatic AP peaks in second derivative was normalized, indicating improved AP initialization and membrane potential acceleration.

### Limitations

While hiPSC-derived neurons from SCA27B patients provide an advanced human model system with the genetic, molecular and cellular background of patients, they present a simplified *in vitro* neuronal model with several limitations. First, the 4–5-week-old stem cell-derived neurons are highly immature [54] and might not adequately mimic the late-onset condition of SCA27B (60.0 years [53.5–68.5], median [IQR]) [54, 55]. Notably, we were able to identify a pronounced phenotype across multiple cellular levels, consistent with findings from FGF14-deficient animal models in which electrophysiological abnormalities precede the onset of measurable motor impairments [8, 13, 19]. Second, despite marked differences in clinical presentation - the monoallelic patient remained clinically episodic, whereas the biallelic patient showed progressive disease burden - we did not observe differences in electrophysiological outcomes or mRNA expression between neurons from the monoallelic and biallelic patient. This finding is noteworthy in light of FA, where GAA•TTC repeat length correlates with frataxin deficiency and several clinical measures. In contrast, for SCA27B exists a comparatively weak genotype–phenotype, which raises the possibility that disease pathogenesis cannot be explained solely by repeat length-dependent FGF14 downregulation and that additional disease-modifying mechanisms may contribute to the observed clinical heterogeneity [56]. At present, our data provide proof of concept that FGF14 transcriptional deficiency is associated with neuronal dysfunction in SCA27B neuronal model, but do not support a straightforward relationship between repeat length, allelic status, and the severity of neuronal dysfunction. Third, the cellular model used in our study does not involve cerebellar but rather cortical glutamatergic neurons, and may thus not well reflect the pathophysiological context of cerebellar disorders. Although FGF14-deficient cerebellar (PC) neurons from rodents pointed into the same direction of a firing deficit [13], validation by future studies in (ideally patient iPSC-derived) PC models is warranted, ideally leveraging cerebellar organoids or animal-derived *in vivo* systems.

## Data availability

Raw data were generated at the *Hertie-Institute for Clinical Brain research, Tübingen University*. Derived data supporting the findings of this study are available from the corresponding author on request.

## Acknowledgements

We gratefully acknowledge Heidi Löffler for her valuable expertise and assistance in generating the electrophysiological data.

